# Ferroptosis mediates the progression of hyperuricemic nephropathy by activating RAGE signaling

**DOI:** 10.1101/2024.05.11.593667

**Authors:** Qiang Wang, Yuemei Xi, Hairong Zhao, De Xie, Linqian Yu, Yunbo Yan, Jiayu Chen, Qian Zhang, Meng Liang, Jidong Cheng

## Abstract

Hyperuricemic nephropathy (HN) represents a prevalent complication arising from hyperuricemia, typified by tubular dysfunction, inflammation, and progressive renal fibrosis, whose pathogenic mechanisms remain enigmatic. Ferroptosis, a newly elucidated iron-dependent form of regulated cell death, plays a role in various disease states. However, its involvement in HN has seldom been explored. In this study, we observed indications of ferroptosis in the renal tissues of urate oxidase knockout (UOX^-/-^) mice, a model of hyperuricemia, as evidenced by increased iron deposition and reduced expression of glutathione peroxidase 4 (GPX4). These findings suggest a substantial role of ferroptosis in the pathogenesis of HN. To further explore this hypothesis, UOX^-/-^ mice were administered Ferrostatin-1, a known inhibitor ferroptosis. This treatment markedly ameliorated tubular injury, necrosis, and inflammatory cell infiltration, mitigated renal fibrosis, reinstated the expression of proteins associated with ferroptosis in renal tissues, and reduced iron overload, lipid peroxidation, and mitochondrial damage in UOX^-/-^ mice. Additionally, we found that receptor for advanced glycation end products (RAGE) propagates ferroptosis-induced renal injury, inflammation and fibrosis, albeit without directly facilitating ferroptosis. Finally, ferroptosis and RAGE upregulation were validated in renal tissues of patients with hyperuricemia-related kidney disease. Collectively, our research elucidates the critical contribution of ferroptosis to HN pathogenesis, indicating that therapeutic strategies targeting ferroptosis and the related RAGE signaling may offer novel therapeutic approaches for managing this condition.

## Introduction

Hyperuricemia has become increasingly prevalent worldwide ^1^. Its overall prevalence has climbed to 14.0% in mainland China ^2^, seriously jeopardizing human health by predisposing to and fueling various metabolic disorders, including gout, hypertension, cardiovascular disease, and chronic kidney disease (CKD) ^3^. However, hyperuricemia often remains asymptomatic during its early stage, resulting in an underestimation of its detrimental impacts by affected individuals.

Responsible for secreting approximately 75% of uric acid, the kidney is one of the main target organs affected by chronic exposure to high levels of uric acid ^4^. Elevated uric acid in the bloodstream exceeds its solubility limit and causes precipitation and deposition of monosodium urate (MSU) crystals within the renal tubules. This cascade of events initiates tubular obstruction, inflammatory responses, and progressive renal fibrosis, eventually culminating in the development of CKD, namely hyperuricemic nephropathy (HN) ^5^. Notably, kidneys manifest merely subtle symptoms until significant functional impairment occurs. Therefore, HN typically has an insidious onset and delayed diagnosis, rendering timely intervention challenging. However, the molecular mechanisms underpinning HN remain poorly understood, highlighting the critical need to elucidate the pathogenic pathways is imperative to identify potential therapeutic targets against HN.

Ferroptosis is a newly recognized form of programmed cell death typified by iron-dependent lipid peroxidation ^6^ ^7^. Among its key regulatory components, system Xc-(catalytic subunit xCT/SLC7A11) and GPX4 are prominent negative factors ^8^. Unlike apoptosis, ferroptosis triggers inflammation by eliciting the release of damage-associated molecular patterns (DAMPs), a shared feature of multiple cell necrosis modalities, including regulated forms like pyroptosis and necroptosis as well as unregulated necrosis ^9^. Emerging evidence implicates ferroptosis in diverse pathological processes, encompassing tumor suppression, ischemia-reperfusion injury, neurodegeneration, acute nephrotoxicity induced by folic acid/cisplatin, and tubular injury in diabetic nephropathy ^10–14^. However, the potential involvement of ferroptosis in HN remains unexplored.

The Receptor for Advanced Glycation End Products (RAGE) belongs to the immunoglobulin superfamily of cell surface molecules. It can be activated by various ligands, including AGE, HMGB1, Decorin, Heme ^15–17^, thereby leading to effects such as systemic insulin resistance, oxidative stress, inflammation, and tissue fibrosis. Under physiological conditions, RAGE is expressed at low levels in renal tubular epithelial cells ^18^. However, during sustained pathological stresses such as diabetic nephropathy or unilateral ureteral obstruction, RAGE can be activated and contribute to renal interstitial fibrosis ^19,20^. Li et al. recently reported that RAGE promotes dysregulation of iron and lipid metabolism in alcoholic liver disease ^21^, while Yang et al. revealed an association between iron overload and RAGE signaling activation in intracerebral hemorrhage ^22^. These findings may imply a potential interplay between RAGE and ferroptosis.

In the present study, we delved into the potential involvement of ferroptosis in HN using urate oxidase knockout (UOX^-/-^) mice as the model of hyperuricemia. We aimed to determine whether targeting ferroptosis and related signaling pathways could confer renoprotection against HN. Our findings may provide new insights into the molecular pathogenesis of HN and identify novel therapeutic strategies.

## Materials and methods

### Reagents

Ferrostatin-1 (#F864515) is from Macklin (China); FPS-ZM1(#HY-19370) is from MedChemExpress (USA); mouse anti-α-SMA (#A2547) antibody is from Sigma (USA); rabbit anti-RAGE (#ab3611), and HMGB1 (#ab18256), as well as mouse anti-TNF-α (#ab255275) antibodies are from Abcam (UK); rabbit anti-E-cadherin (#CPA1199) antibody is from Cohesion Biosciences (UK); rabbit anti-HO-1 (#10701-1-AP) antibody is from Proteintech (USA); rabbit anti-Iba1 (#019-19741) is from Wako (Japan); rabbit anti-LC3B (#3868), Phospho (p)-P38 MAPK (Thr180/Tyr182) (#4511), P38 MAPK (#8690), p-JNK (Thr183/Tyr185) (#4668), JNK (#9252), p-ERK1/2 (Thr202/Tyr204) (#4370), ERK (#4695), p-NF-κB p65 (Ser536) (#3033), NF-κB p65 (#3034) antibodies are from Cell Signaling Technology (USA); mouse anti-GAPDH (#AC033), rabbit anti-β-actin (#AC026), xCT (#A2413), GPX4 (#A13309, #A11243), TfR1(#A4865), COX2 (#A3560), P62 (#A19700), 4-HNE (#A2245), HRP-conjugated goat anti-rabbit IgG (#AS014), HRP-conjugated goat anti-mouse IgG (#AS003) antibodies are from Abclonal (China). Enhanced Chemiluminescent (ECL) Substrate (#BMU102-CN) is from Abbkine (China). Kits for determining biochemical indexes of uric acid (#C012-2-1) and creatinine (#C011-2-1), and urea nitrogen (#C013-2-1 or #BC1535) are from Nanjing Jiancheng (China) or Solarbio (China).

### Mice

Due to the presence of urate oxidase (UOX), mice can further convert uric acid into the more soluble allantoin. Thus, pharmacological inhibition or genetic knockout of UOX are common strategies to establish hyperuricemic mouse models ^23^. We previously generated a hyperuricemic model by knocking out UOX gene in C57BL/6J mice, namely, UOX^-/-^ mice ^24^. They exhibited significant hyperuricemia, insulin resistance, kidney injury and hepatic fat accumulation ^24,25^. Starting at six weeks of age, mice were administered with corresponding drugs (ferroptosis inhibitor Fer-1, RAGE inhibitor FPS-ZM1) for 1 month before euthanasia. Fer-1 (1% DMSO in normal saline) was intraperitoneally administered at 2 mg/kg per day. FPS-ZM1 (1% DMSO in normal saline) was also delivered via intraperitoneal injection at 1.5 mg/kg per day. At the end of the treatment period, mice were anaesthetized with isoflurane, and then sacrificed by cervical dislocation. All mice were housed in the Laboratory Animal Center of Xiamen University, having free access to food and water. All animal experiments were revised and approved by the Animal Ethics Committee of Xiamen University (approval number: XMULAC20200122).

### Human Renal Biopsy Specimens

The specimens of hyperuricemia-associated nephropathy were derived from renal biopsy. Control samples were derived from healthy adjacent non-cancerous tissues of individuals undergoing tumor nephrectomies without diabetes or chronic kidney disease. Each group includes two samples. The acquisition and utilization of tissue specimens were reviewed and approved by the Medical Ethics Committee of Xiang’an Hospital, Xiamen University (approval number: XAHLL2023014).

### Preparation of MSU crystals

Uric acid (750 mg) was dissolved in a 0.5 M sodium hydroxide solution while undergoing continuous stirring to achieve a 15 mg/mL urate solution. When left undisturbed at 4°C overnight, visible precipitates formed. The solution was then allowed to continue incubating at 4°C, enabling a gradual crystallization of MSU crystals, characterized by needle-like morphology under microscope. These crystals were collected through a second centrifugation step, followed by a thorough drying and subsequent weighing. To ensure sterility, the MSU crystals underwent autoclaving before being dissolved in PBS to yield a stock solution of 20 mg/mL.

### Cell culture

The human kidney 2 (HK2) cell line, derived from human normal kidney, is obtained from Center for Excellence in Molecular and Cellular Sciences, Chinese Academy of Sciences (China), routinely cultured with DMEM/F-12 (#L310KJ, BasalMedia, China) containing 10% fetal bovine serum in a 37℃incubator with 5% CO_2_.

### Cell Counting Kit-8 (CCK8) Assay

CCK8 assay was used to assess cell viability as per manufacturer’s instructions. Briefly, 5000 cells per well were seeded into 96-well plates, treated with corresponding compounds, and then incubated for indicated time. The old medium was replaced with complete medium containing 10% CCK8 reagent and further incubated for 1.5h before measuring the absorbance at 450 nm using a microplate reader (#Multiskan Sky, Thermo, USA). Cell viability was calculated as a percentage relative to untreated controls.

### Lactate dehydrogenase (LDH) release assay

Cells were seeded in 6 cm dishes, treated with appropriate reagents or a control vehicle. A blank control group with cell-free complete medium was set. Each dish contained 3 ml of culture medium. After 24 hours, the culture medium from each dish was centrifuged to obtain 100 µl of the supernatant. Subsequently, the cells underwent two washes with PBS, followed by the addition of 3 ml of 0.2% Triton X-100 to fully lyse the cells, after which 100 µl of the lysate supernatant was collected. The LDH content in both the cell supernatant and the cell lysate was determined using a LDH assay kit (#C0016, Beyotime, China). The percentage of LDH release was calculated using the formula: [LDH in the supernatant / (LDH in the supernatant + LDH in the cell lysate)] * 100%.

### Biochemical Analysis

The serum levels of uric acid (UA), creatinine (CREA), and blood urea nitrogen (BUN) were determined in accordance with the respective manufacturer’s protocols. Briefly, serum samples were allowed to react with the specific reaction mixtures tailored to each biochemical index for the indicated duration. The optical density at the specific wavelength was recorded using a microplate reader.

### Histological Analysis and tubular damage evaluation

Fresh mouse kidney specimens were fixed by perfusion and immersion with 4% paraformaldehyde solution. Subsequently, the tissues were embedded in paraffin and sectioned to a thickness of 5 μm. These sections underwent the standard deparaffinization and rehydration procedures and then were subject to staining with hematoxylin and eosin (H&E), periodic acid-Schiff (PAS), and Masson’s trichrome according to established protocols. Following staining, the sections were dehydrated, cleared, and mounted with a resinous mounting medium. Tubular damage in sections stained with H&E was assessed based on the extent of the damaged area, using the following scale: 0 = normal, 1 = 1-25%, 2 = 26-50%, 3 = 51-75%, and 4 = 76-100%.

### Immunohistochemistry

Paraffin-embedded tissue sections were deparaffinized, rehydrated, and then subjected to heat-mediated antigen retrieval in citrate buffer (pH 6.0). To quench endogenous peroxidase activity the sections were treated with a 3% hydrogen peroxide solution. Subsequently, the sections were blocked using 5% normal goat serum and then incubated with rabbit anti-Iba1 (1:200), GPX4 (1:100) or RAGE (1:100) antibodies overnight at 4°C. After washing, the sections were exposed to HRP-conjugated secondary antibody for 1 hour at room temperature. Immunostaining was developed using the DAB Peroxidase Substrate Kit (#DAB-0031, MXB biotechnology, China), and the sections were counterstained with hematoxylin. Images were captured using a light microscope.

### Immunofluorescence

Paraffin-embedded tissue sections were deparaffinized, rehydrated, and then subjected to heat-mediated antigen retrieval in citrate buffer (pH 6.0). After blocking with 5% normal goat serum, the sections were incubated with rabbit anti-GPX4 (1:100) antibody overnight at 4°C. After washing, the sections were incubated with Alexa Fluor-conjugated secondary antibody for 1 hour at room temperature. Nuclei were counterstained with DAPI, and fluorescent images were acquired using a fluorescence microscope (#DM2700 P, Leica, Germany).

### Iron deposition evaluation with ferrous iron colorimetric assay and DAB-enhanced Prussian blue staining

For ferrous iron measurement, fresh kidney tissues (∼30 mg) were homogenized in 300 μL of a specified extraction solution, followed by centrifugation to yield a clear supernatant. An aliquot of 200 μL from each sample or and iron standard was incubated with 150 μL of Chromogenic Solution at 37°C for 10 min in 1.5 mL microcentrifuge tubes. After incubation, 200 μL of the supernatant was transferred into the corresponding wells of a 96-well microplate. The OD values were measured at 593 nm using a microplate reader. The ferrous iron concentration in the samples was calculated by comparing their OD values against that of the iron standard.

For tissue iron visualization, DAB-enhanced Prussian blue staining was conducted. Briefly, kidney sections were incubated with Prussian blue staining solution for 20 min at 37℃, followed by exposure to a DAB substrate solution for 10 min at 37℃. After counterstaining with hematoxylin, the sections were dehydrated, cleared and mounted. Images were acquired using a light microscope.

### Transmission Electron Microscopy (TEM)

To prepare samples for TEM, we harvested the upper pole renal cortex of mice and fixed them in 2.5% neutral glutaraldehyde and 1% osmium acid at 4°C overnight. Subsequently, the tissues were dehydrated, embedded, and cut into 50 nm ultrathin sections, which were then collected on copper grids. For enhanced contrast, double staining was performed using uranyl acetate and lead nitrate. Finally, tubular mitochondria were visualized and recorded using a Hitachi HT-7800 TEM (Japan).

### Quantitative PCR (qPCR)

Total RNA was extracted from kidney cortices or cells using RNA extract solution (#G3010, Servicebio, China) following the manufacturer’s instructions. Subsequently, 2 μg of RNA were subject to reverse transcription using an Evo M-MLV Mix Kit (#AG11728, Accurate Biotechnology, China). qPCR was conducted on a Real-Time System (#CFX96, Bio-Rad, USA) using SYBR Green mix (#RK21219, Abclonal, China). ACTB or GAPDH was used as the internal control, and relative mRNA levels were calculated using the 2−ΔΔCt method. The primer sequences are listed in Table S1.

### Detection of Lipid Peroxidation

For cultured live cells, lipid peroxidation was evaluated using the fluorescent probe C11 BODIPY 581/591 (#RM02821, Abclonal, China). In brief, cells seeded on coverslips in 12-well plates were allowed to adhere overnight. They were then treated with MSU crystals for 48 h and subsequently incubated with 5 μM C11 BODIPY 581/591 for another 1 hour at 37°C. Following a PBS wash, the coverslips were inverted onto glass slides and imaged using a fluorescence microscope (#DM2700 P, Leica). Both red (reduced) and green (oxidized) fluorescence images were captured.

For kidney sections, 4-HNE was selected as the marker of lipid peroxidation by immunofluorescence.

For fresh kidney samples, MDA levels were used as an indicator of lipid peroxidation. Briefly, kidney tissues underwent homogenization followed by centrifugation to extract the supernatant. An MDA assay kit (#A003-1, Nanjingjiancheng, China) was used according to the manufacturer’s guidelines. To ensure consistent comparative analysis, the obtained MDA content was normalized against the weight of the kidney tissue.

### Intracellular Ferrous Iron Detection Using FerroOrange

Intracellular ferrous iron levels were determined using FerroOrange (#F374, Dojindo, Japan). Briefly, cells were seeded in a 6-well plate and allowed to adhere overnight. After drug intervention, cells were washed three times with serum-free culture medium, followed by the addition of 1 µM FerroOrange. Following a 30-min incubation at 37°C, the cells were observed and photographed under a fluorescence microscope. The red fluorescence represents the relative ferrous iron content.

### Western blot (WB) analysis

WB analysis was performed as previously described ^26^. Briefly, cells or tissues were lysed in RIPA buffer containing protease and phosphatase inhibitors. Equal amounts of protein lysates were separated by SDS-PAGE and subsequently transferred to PVDF membranes. These membranes were blocked with 5% non-fat milk at room temperature for 30 min, incubated with primary antibodies overnight at 4°C, and then with HRP-conjugated secondary antibodies at room temperature for 1.5 h. Protein bands were visualized using ECL substrate in a chemiluminescence imaging system (#ChemiScope 6100, CLiNX, China) and band intensities were quantitated by densitometry using ImageJ software (NIH, USA).

### Enzyme-Linked Immunosorbent Assay (ELISA)

The levels of HMGB1 protein in cell supernatants and serum were measured according to the manufacturer’s instructions (#E-EL-H1554, #E-EL-M0676, Elabscience, China). Briefly, a complex of anti-HMGB1 coat antibody, HMGB1 present in supernatants or serum, biotinylated anti-HMGB1 detection antibody, and HRP-conjugated streptavidin was formed, followed by reaction with TMB substrate, which was terminated by the addition of stop solution and absorbance was measured at 450 nm. HMGB1 concentration was calculated from a standard curve.

### Statistical analysis

Statistical analyses were performed using GraphPad Prism 9 software. Differences between two groups were analyzed by Student’s t-test. One-way ANOVA followed by Tukey’s multiple comparison test was used for multiple comparisons. The data were present as mean ± standard error of mean (S.E.M). *P* < 0.05 was considered statistically significant.

## Results

### 1. Occurrence of ferroptosis in the kidney of UOX^-/-^ mice

By the age of eight weeks, UOX^-/-^ mice displayed notably elevated serum levels of UA, CREA, and BUN compared to their wild-type (WT) counterparts (Fig. 1A), indicating renal impairment in the HN model. PAS staining demonstrated extensive tubular casts and necrosis in the kidneys of UOX^-/-^ mice, further corroborating renal injury (Fig. 1C). WB analysis of renal cortex lysates showed marked upregulation of α-SMA, a pivotal mesenchymal marker, in UOX^-/-^ mice (Fig. 1B). These findings imply that increased levels of serum UA induced renal damage, ultimately leading to kidney fibrosis in UOX^-/-^ mice, which was confirmed by MASSON staining (Fig. 1C). Furthermore, significant downregulation of the key anti-ferroptosis markers, GPX4 and xCT, was observed in the kidneys of UOX-/- mice (Fig. 1B), alongside with an increased iron deposition indicated by DAB-enhanced Prussian blue staining (Fig. 1C). Immunohistochemistry (IHC) showed that GPX4 was predominantly expressed in renal tubules instead of glomeruli, and its expression was diminished in UOX^-/-^ mouse kidneys (Fig. 1C). Collectively, these results implied the occurrence of ferroptosis in HN.

**Figure 1.**
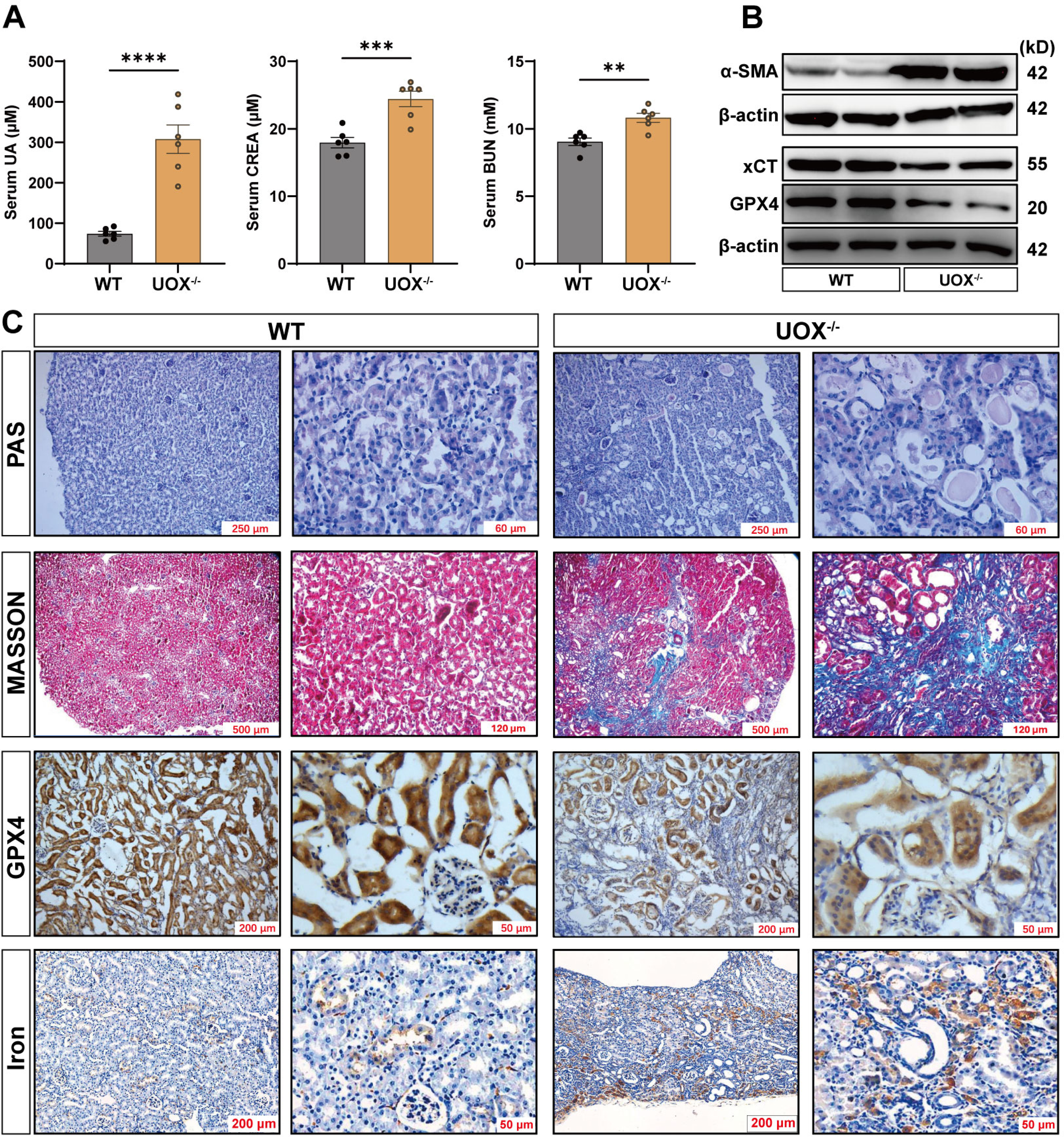
Renal tissues from UOX^-/-^ mice exhibit ferroptosis-related changes. (A) Increased serum UA, CREA and BUN detected by biochemical assay in mice. Data are presented as mean ± S.E.M (n = 6). (B) Renal protein levels of α-SMA, GPX4 and xCT detected by WB analysis in mice. (C) Renal staining in mice included PAS, Masson, IHC against GPX4, and DAB-enhanced Prussian blue staining. WT, wild type; UOX^-/-^, urate oxidase knockout; IHC, immunohistochemistry; UA, uric acid; CREA, creatinine; BUN, blood urea nitrogen. Scale bars are marked in figures. \*\**P* < 0.01, \*\*\**P* < 0.001, \*\*\*\**P* < 0.0001.

### 2. Ferroptosis inhibitor Fer-1 ameliorates renal injury and ferroptosis in UOX^-/-^ mice

Mice received either vehicle or Fer-1 treatment for one month. The results indicated that in UOX^-/-^ mice, Fer-1 substantially reduced elevated serum BUN and CREA levels without altering serum UA levels (Fig. 2A-C). qPCR analysis revealed that Fer-1 significantly decreased mRNA levels of renal injury markers KIM1 and LCN2 in UOX^-/-^ mouse kidney (Fig. 2D-E). H&E staining revealed remarkable tubular dilation, necrosis and immune cell infiltration in UOX^-/-^ mouse kidney, all of which showed marked improvement upon Fer-1 treatment (Fig. 2F-G). The indicators of lipid peroxidation (4-HNE and MDA) were significantly upregulated in the kidneys of UOX-deficient mice, whereas Fer-1 treatment effectively inhibited their elevation (Fig. 2H-I). Furthermore, we scrutinized the expression of ferroptosis-related markers and noticed that Fer-1 administration restored the diminished protein levels of key ferroptosis markers TfR1, xCT, and GPX4 in the kidneys of UOX^-/-^ mice (Fig. 2J-K). The restorative effect of Fer-1 on GPX4 expression was corroborated by immunofluorescence (Fig. 2M). Furthermore, we observed iron deposition by ferrous iron colorimetric assay and DAB-enhanced Prussian blue staining as well as mitochondrial shrinking with disappearance of mitochondrial cristae by TEM in renal tubular epithelial cells of UOX^-/-^ mice, which were effectively reversed by Fer-1 treatment (Fig. 2L, N-O). Additionally, we determined the mRNA levels of regulatory genes involved in ferroptosis (DMT1, IRP1, IRP2, FPN1, FTL1, FTH1, etc.) to further elucidate the role of ferroptosis in HN (Fig. 2P).

**Figure 2.**
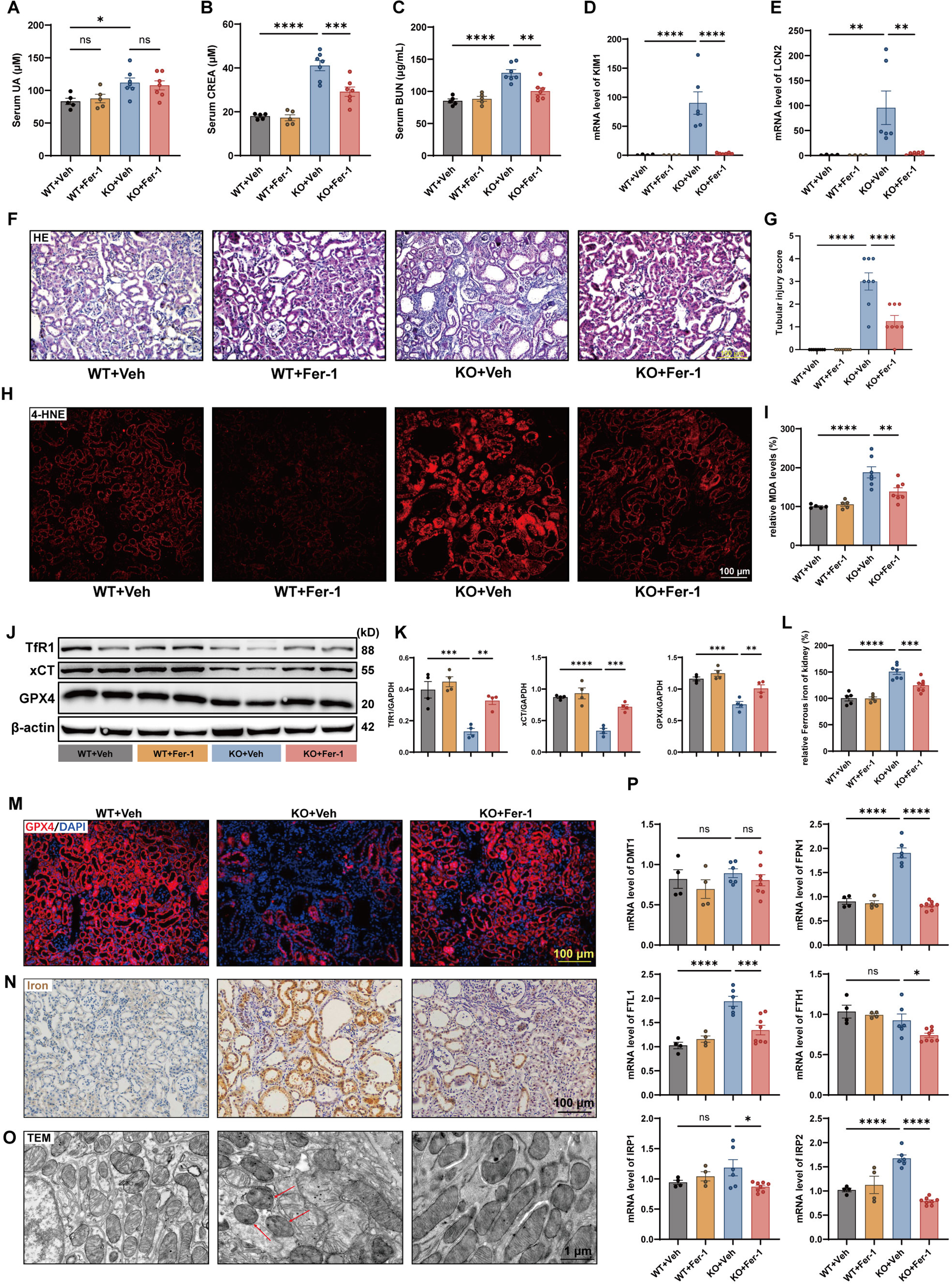
Ferroptosis inhibitor Fer-1 ameliorates renal injury and ferroptosis in UOX^-/-^ mice. Serum levels of UA (A), CREA (B), and BUN (C). mRNA levels of renal injury markers KIM1 (D) and LCN2 (E) determined by qPCR. (F) H&E staining of mouse kidney and the corresponding tubular injury score (G). Scale bar = 100 μm. (H) Immunofluorescence of the lipid peroxidation marker 4-HNE in kidney tissue sections. Scale bar = 100 μm. (I) Relative content of the lipid peroxidation marker MDA in kidney homogenates. (J) Protein levels of TfR1, xCT and GPX4 in mouse kidney detected by WB analysis and their semi-quantification (K). (M) GPX4 shown by immunofluorescence and cell nucleus indicated by DAPI in mouse kidney. Scale bar = 100 μm. Iron deposition indicated by ferrous iron colorimetric assay (L) and DAB-enhanced Prussian blue staining (N) in mouse kidney. Scale bar = 100 μm. (O) Transmission electron microscopy revealed the mitochondrial morphology of mouse renal tubular epithelial cells. Scale bar = 1 μm. (P) mRNA levels of regulatory genes of ferroptosis detected by qPCR in mouse kidney. Data are presented as mean ± S.E.M (n = 5 or 7 for A-C, n = 4∼8 for D-E, n = 8 for G, n = 5 or 7 for I, n = 4 for J-K, n = 4∼8 for P). KO, UOX^-/-^ knockout, Veh, vehicle, Fer-1, Ferrostatin-1. \**P* < 0.05, \*\**P* < 0.01, \*\*\**P* < 0.001, \*\*\*\**P* < 0.0001.

Collectively, renal injury in hyperuricemic UOX^-/-^ mice was mediated by ferroptosis, and the amelioration of renal injury in UOX^-/-^ mice was achieved through the inhibition of ferroptosis.

### 3. Ferroptosis inhibitor Fer-1 alleviates renal inflammation in UOX^-/-^ mice

Considering the immunogenic nature of ferroptosis, we performed IHC staining for Iba1, a marker of peripheral macrophages, to assess macrophage infiltration in the kidney. Extensive macrophage infiltration was observed in the tubulointerstitium of the UOX^-/-^ mouse kidney, which was significantly reduced by Fer-1 treatment (Fig. 3A). Furthermore, qPCR of the renal cortex showed increased mRNA levels of inflammation-related markers (IL1β, IL6, TNF-α, COX2, MCP-1) in UOX^-/-^ mice, all of which were notably suppressed by Fer-1 to levels comparable to those of WT mice (Fig. 3B). We further examined expression levels of the inflammation-related proteins COX2, TNF-α, p-P65, HMGB1, and MAPK family members (ERK, P38 MAPK, JNK), which possess vital functions in renal injury, by WB analysis. Compared to WT mice, their expression levels were significantly increased in the UOX^-/-^ mouse kidney, whereas Fer-1 markedly counteracted these changes (Fig. 3C-F). Notably, COX2 has been identified as an important ferroptosis marker in recent years ^27^, and HMGB1 plays a key role in ferroptosis-induced inflammation ^28^. Since HMGB1 is an important DAMP released by ferroptotic cells, we further measured serum HMGB1 levels by ELISA. The results indicated that serum HMGB1 levels followed a comparable but notably more pronounced expression pattern compared with renal HMGB1 protein levels. Serum HMGB1 significantly surged in UOX^-/-^ mice, which was markedly inhibited by Fer-1 treatment (Fig. 3G). Taken together, ferroptosis mediated the inflammatory response in the UOX^-/-^ mouse kidney.

**Figure 3.**
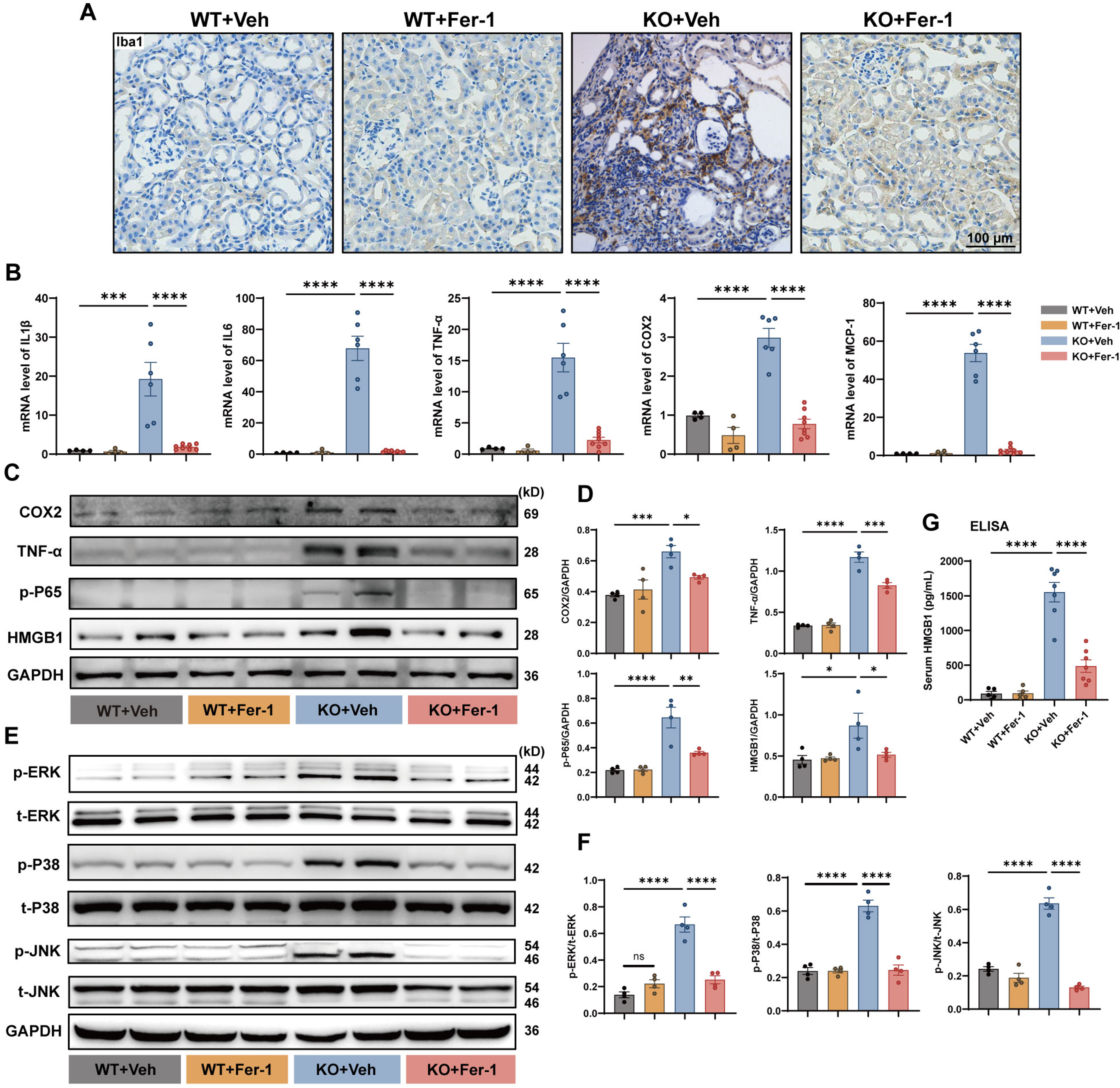
Ferroptosis inhibitor Fer-1 alleviates renal inflammation in UOX^-/-^ mice. (A) Macrophage infiltration shown by IHC staining in mouse kidney. Scale bar = 100 μm. (B) mRNA expression of inflammation-related indicators, including pro-inflammatory factors IL1β, IL6 and TNF-α, as well as chemokines COX2 and MCP-1 determined by qPCR in mouse kidney. (C) Protein levels of COX2, TNF-α, p-P65, HMGB1 determined by WB analysis and their semi-quantification (D). (E) Protein levels of the MAPK family (ERK, P38 MAPK, JNK and their phosphorylated counterparts) and determined by WB analysis and their semi-quantification (F). (G) Serum levels of HMGB1 determined by ELISA. Data are presented as mean ± S.E.M (n = 4∼8 for B, n = 4 for C-F, n = 4 for F, n = 4∼8 for G). ns, not significant, \**P* < 0.05, \*\*\**P* < 0.001, \*\*\*\**P* < 0.0001.

### 4. Ferroptosis inhibitor Fer-1 mitigates renal autophagy and fibrosis in UOX^-/-^ mice

Autophagy has been found to induce inflammatory injury and fibrosis in HN ^29^, and it’s also worth noting that ferroptosis is a process that depends on autophagy ^30^. In the UOX^-/-^ mouse kidney, we observed an increase of LC3B □/□ ratio and a concurrent decrease of P62, the two key markers of autophagy, which indicate sustained autophagic activation. Notably, inhibiting ferroptosis with Fer-1 significantly attenuated autophagy (Fig. 4A). In terms of renal fibrosis, the pivotal epithelial marker E-cadherin was downregulated while α-SMA was upregulated in UOX^-/-^ mice, highlighting remarkable renal fibrosis. Fer-1 markedly curtailed renal fibrosis in UOX^-/-^ mouse (Fig. 4A-B). Masson staining confirmed the significant tubulointerstitial fibrosis in UOX^-/-^ mouse kidney, which was markedly inhibited by Fer-1 (Fig. 4C-D).

**Figure 4.**
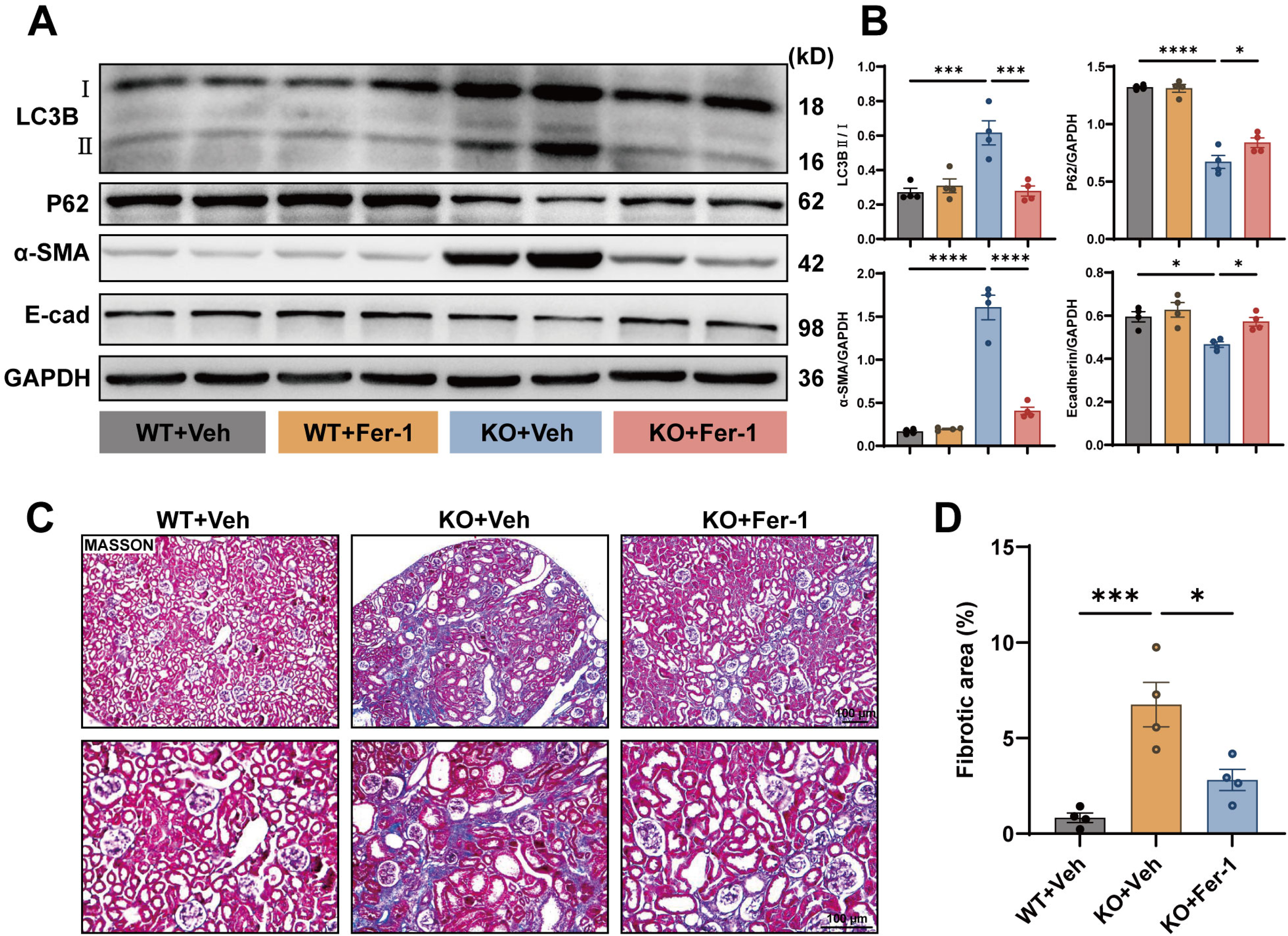
Ferroptosis inhibitor Fer-1 mitigates renal autophagy and fibrosis in UOX^-/-^ mice. (A) Protein levels of LC3B, P62, E-cadherin and α-SMA detected by WB in mouse kidney and their semi-quantification (B). (C) Renal fibrosis shown by Masson staining and its semi-quantification (D). Data are presented as mean ± S.E.M (n = 4). Scale bar = 100 μm. \**P* < 0.05, \*\*\**P* < 0.001, \*\*\*\**P* < 0.0001.

Taken together, ferroptosis inhibition significantly halts renal autophagy and subsequent fibrosis in UOX^-/-^ mice.

### 5. MSU crystals induce ferroptosis in HK2 cells

Through WB analysis, we showed that MSU crystals led to a significant decrease in the protein levels of GPX4 and xCT in HK2 cells. Meanwhile, fibrogenesis was boosted as evidenced by the upregulation of α-SMA and the downregulation of E-cadherin (Fig. 5A). Additionally, MSU crystal treatment significantly enhanced lipid peroxidation (green fluorescence) in HK2 cells by using the fluorescent probe C11 BODIPY 581/591, while Fer-1 inhibited this change (Fig. 5B). For MAPK signaling, our findings revealed that inhibiting ferroptosis terminated the activation of p-ERK without discernible effect on the p-P38 MAPK and p-JNK (Fig. 5C). Additionally, we observed that inhibiting ferroptosis mitigated cellular autophagy induced by MSU crystals (Fig. 5C). By using CCK8 and LDH release assays, we demonstrated that Fer-1 partially restored HK2 cell injury (Fig. 5D-E). Next, we measured HMGB1 levels in the supernatants of HK2 cells. The results revealed that MSU crystals significantly induced the release of HMGB1, a process that was effectively halted by the inhibition of ferroptosis with Fer-1 (Fig. 5F). FerroOrange was employed to label intracellular ferrous iron. The findings revealed that MSU crystals induced an accumulation of intracellular ferrous iron, which was mitigated by inhibiting ferroptosis (Fig. 5G). Collectively, ferroptosis partially mediated MSU crystals-induced injury in renal tubular epithelial cells.

**Figure 5.**
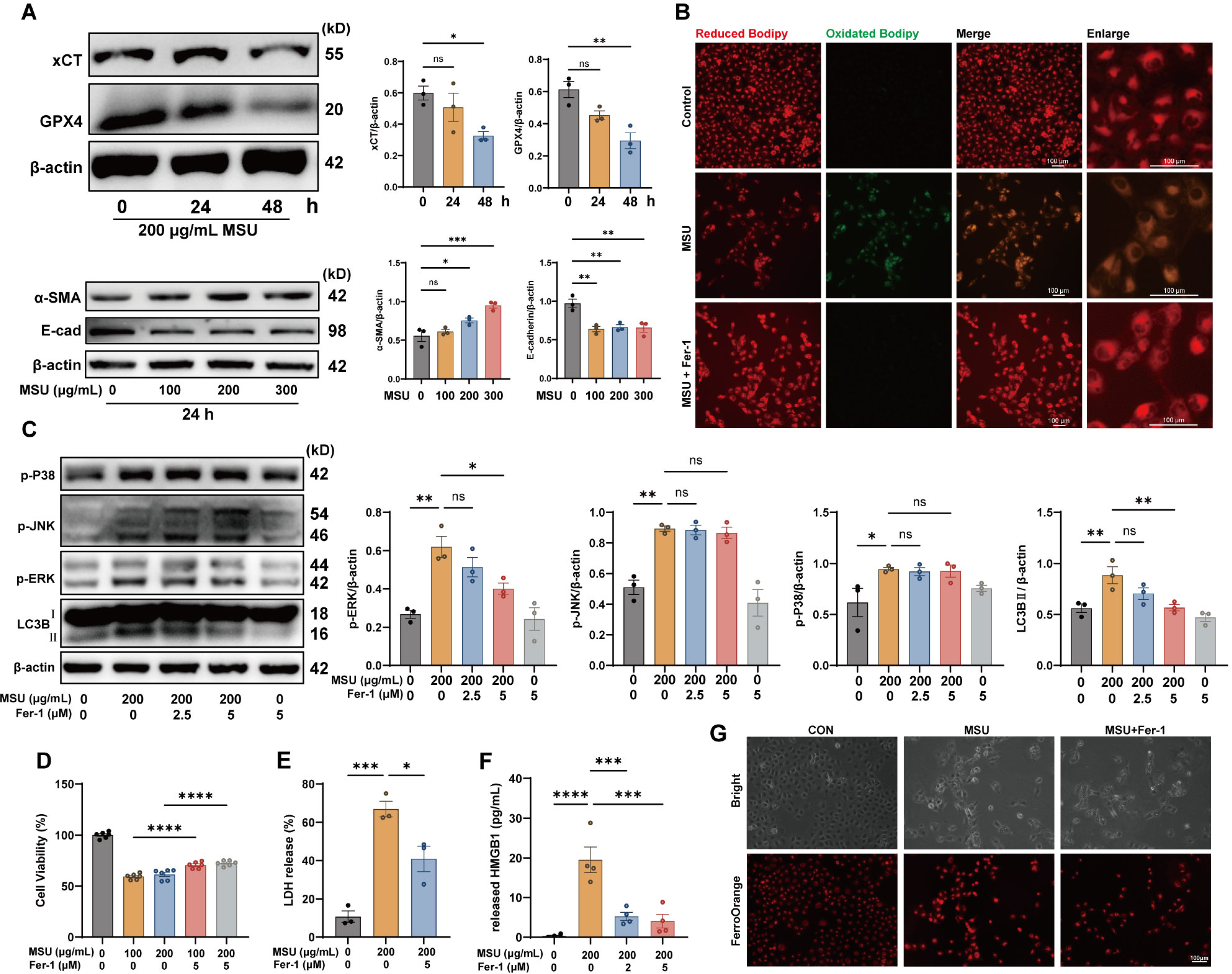
MSU crystals induce ferroptosis in HK2 cells. (A) Protein levels of GPX4, xCT, α-SMA, E-cadherin detected by WB in HK2 cells. β-actin serves as the internal reference protein. (B) Detection of lipid peroxidation in HK2 cells using C11-BODIPY 581/591 fluorescent probe. HK2 cells were subject to treatment with a vehicle (Control), 200 μg/mL MSU crystals (MSU), or a combination of 200 μg/mL MSU crystals and 5 μM Fer-1 (MSU+Fer-1) for 48 h. The red and green fluorescence represent the reduced and oxidized forms, respectively. The transition from red fluorescence to green fluorescence signifies the presence of lipid peroxidation. Scale bar = 100 μm. (C) Protein levels of MAPK family (p-P38 MAPK, p-JNK, p-ERK) and LC3B detected by WB in HK2 cells. β-actin serves as the internal reference protein. HK2 cell injury evaluated by CCK8 assay (D) and LDH release assay (E). (F) HMGB1 release detected by ELISA in HK2 cells. (G) Intracellular ferrous iron indicated by FerroOrange. Data are presented as mean ± S.E.M (n = 3 for A, C and E, n = 6 for D, n = 4 for F). Scale bar = 100 μm.\*\*\**P* < 0.001, \*\*\*\**P* < 0.0001.

### 6. RAGE inhibition alleviates renal injury induced by ferroptosis in UOX^-/-^ mice

qPCR, WB and IHC analysis consistently unveiled that Fer-1 administration curbed RAGE upregulation in UOX^-/-^ mouse kidney (Fig. 6A-C), indicating that RAGE signaling is activated in response to ferroptosis in HN. Therefore, we speculated that RAGE might play an important role in HN. In light of this, a comprehensive exploration was undertaken to ascertain whether RAGE mediates ferroptosis, inflammation, or fibrosis in HN.

**Figure 6.**
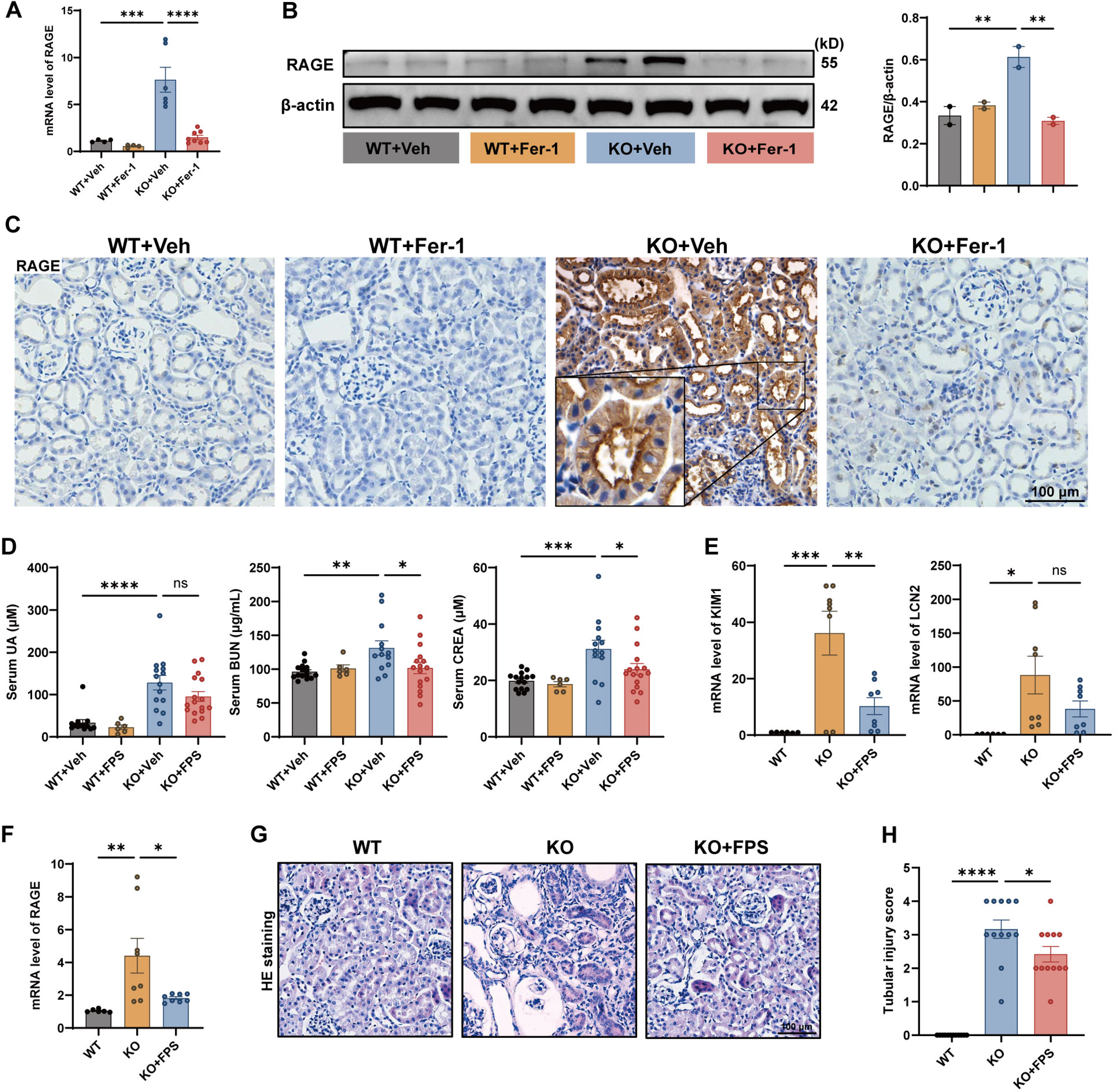
Pharmacological inhibition of RAGE ameliorates renal injury in UOX^-/-^ mice. (A-C) RAGE expression detected by qPCR, WB and IHC in mouse kidney. (D) Serum levels of UA, BUN, CREA. (E) mRNA levels of renal injury markers KIM1 and LCN2 detected by qPCR in mice kidney. (F) mRNA levels of RAGE detected by qPCR in mice kidney. (G) Tubular injury shown by H&E staining in mice kidney. Data are presented as mean ± S.E.M (n = 4∼8 for A, n = 2 for B, n = 6∼16 for D, n = 6∼8 for E and F, n = 12 for H). Scale bar = 100 μm. ns, not significant, \**P* < 0.05, \*\**P* < 0.01, \*\*\**P* < 0.001, \*\*\*\**P* < 0.0001.

Mice were treated with either vehicle or a specific RAGE inhibitor FPS-ZM1. The results showed that FPS-ZM1 significantly suppressed the increased serum CREA and BUN levels in UOX^-/-^ mice without altering the elevated serum UA levels (Fig. 6D). In the renal cortex, qPCR revealed a remarkable attenuation of the overexpressed renal injury markers, namely KIM1 and LCN2, following FPS-ZM1 administration (Fig. 6E). Furthermore, the inhibitory effect of FPS-ZM1 on RAGE expression was confirmed through qPCR analysis (Fig. 6F). H&E staining revealed that FPS-ZM1 intervention ameliorated tubular injury in UOX^-/-^ mouse kidney (Fig. 6G-H). Nevertheless, FPS-ZM1 failed to reinstate the diminished protein levels of core ferroptosis regulatory proteins, GPX4 and xCT (Fig. S1A). What’s more, RAGE inhibition did not alleviate renal iron deposition in UOX^-/-^ mice (Fig. S1B). These data imply that inhibiting RAGE does not impede ferroptosis. Collectively, ferroptosis in HN condition activates RAGE, and RAGE inhibition alleviates renal injury induced by ferroptosis.

### 7. RAGE mediates ferroptosis-induced renal inflammation in UOX^-/-^ mice

IHC staining delineated extensive infiltration of macrophage (labeled by Iba1) in UOX^-/-^ mouse kidney, particularly surrounding necrotic tubules. However, FPS-ZM1 conspicuously curtailed the extent of macrophage infiltration (Fig. 7A). We further examined the expression profile of inflammation-related proteins. Compared to WT mice, UOX^-/-^ mouse kidney showed remarkably increased protein levels of TNF-α, COX2, HMGB1, and an increasing tendency for RAGE and oxidative stress-responsive protein HO-1, underscoring a pronounced presence of inflammation and oxidative stress. Intriguingly, UOX^-/-^ mice subject to FPS-ZM1 intervention exhibited marked abrogation of the aforementioned alterations (Fig. 7B-C). qPCR results showed that FPS-ZM1 markedly inhibited the upregulated mRNA levels of inflammation-related genes (MCP-1, TNF-α, IL1β, IL6). However, FPS-ZM1 did not significantly affect the mRNA levels of COX2 and HMGB1 (Fig. 7D). This suggests that FPS-ZM1 may potentially inhibit COX2 at the post-transcriptional level, while mainly suppressing HMGB1 secretion rather than transcription.

**Figure 7.**
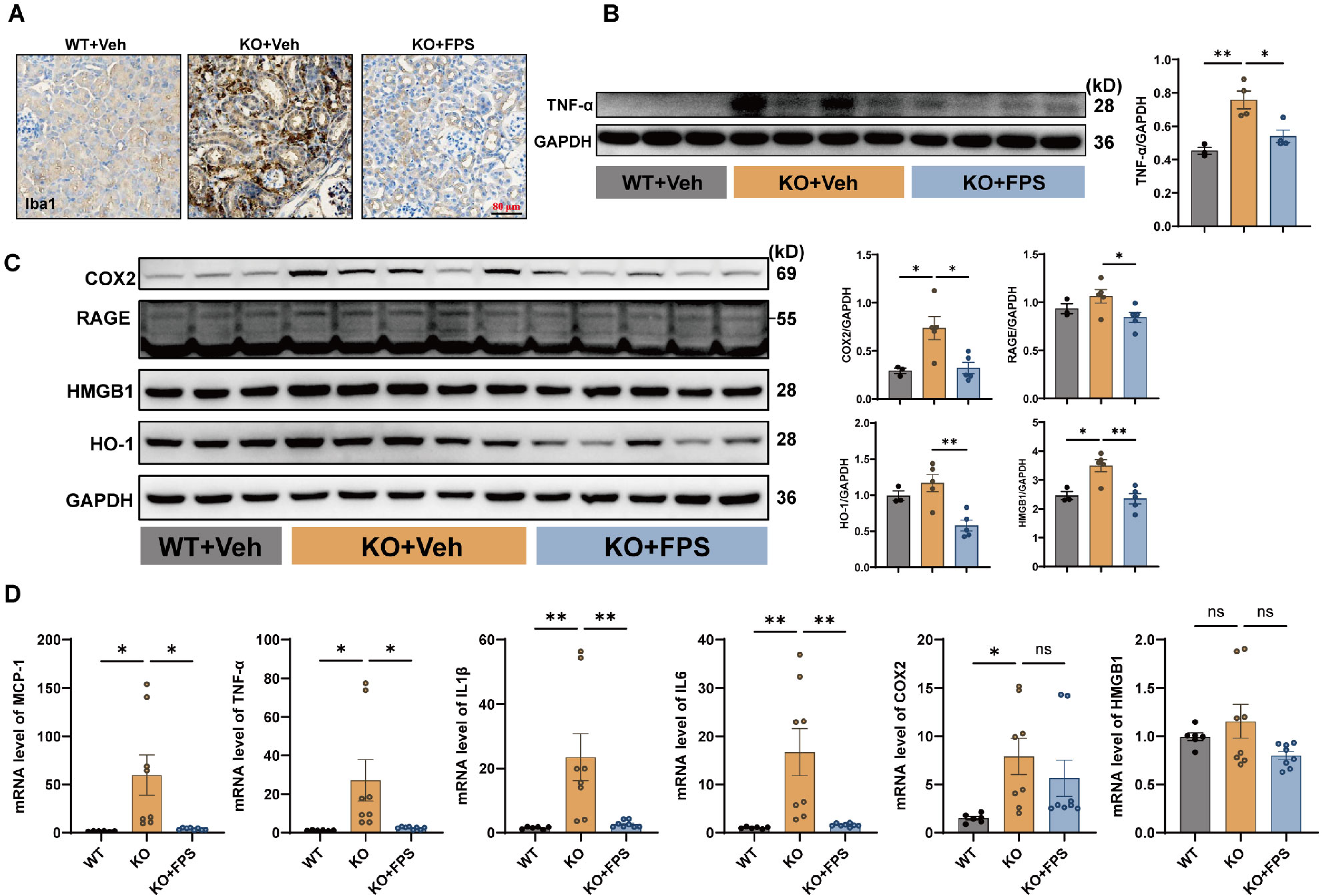
Pharmacological inhibition of RAGE alleviates inflammation in UOX^-/-^ mouse kidney. (A) Macrophage infiltration shown by IHC in mouse kidney. Scale bar = 50 μm. (B, C) Protein levels of TNF-α, COX2, RAGE, HMGB1, HO-1in mouse kidney detected by WB analysis and its semi-quantification. GAPDH serves as the internal reference protein. (D) mRNA levels of MCP-1, TNF-α, IL1β, IL6, COX2, HMGB1 determined by qPCR. Data are presented as mean ± S.E.M (n = 3∼4 for B, n = 3∼5 for C, n = 6∼8 for D). FPS, FPS-ZM1. ns, not significant, \**P* < 0.05, \*\**P* < 0.01.

### 8. RAGE inhibition mitigates renal autophagy and fibrosis in UOX^-/-^ mice

In congruence with the forementioned results, the LC3BII/I ratio was significantly increased in UOX^-/-^ mouse kidney, indicating an enhanced autophagic process, which was significantly abrogated by FPS-ZM1 administration (Fig. 8A-B). Similarly, the expression patterns of E-cadherin and α-SMA indicated that FPS-ZM1 markedly reversed renal fibrosis in UOX^-/-^ mice (Fig. 8C-D). Notably, p-ERK, a crucial co-factor in RAGE-driven kidney fibrosis ^31^, was significantly inhibited in response to FPS-ZM1 treatment. Masson staining further morphologically conformed that FPS-ZM1 inhibited tubulointerstitial fibrosis in UOX^-/-^ mouse kidney (Fig. 8E). In HK2 cells, RAGE inhibition with FPS-ZM1 suppressed MSU crystals-induced p-ERK, p-P38 MAPK, COX2, autophagy, α-SMA, while its influence on p-JNK remained negligible (Fig. S2A-D).

**Figure 8.**
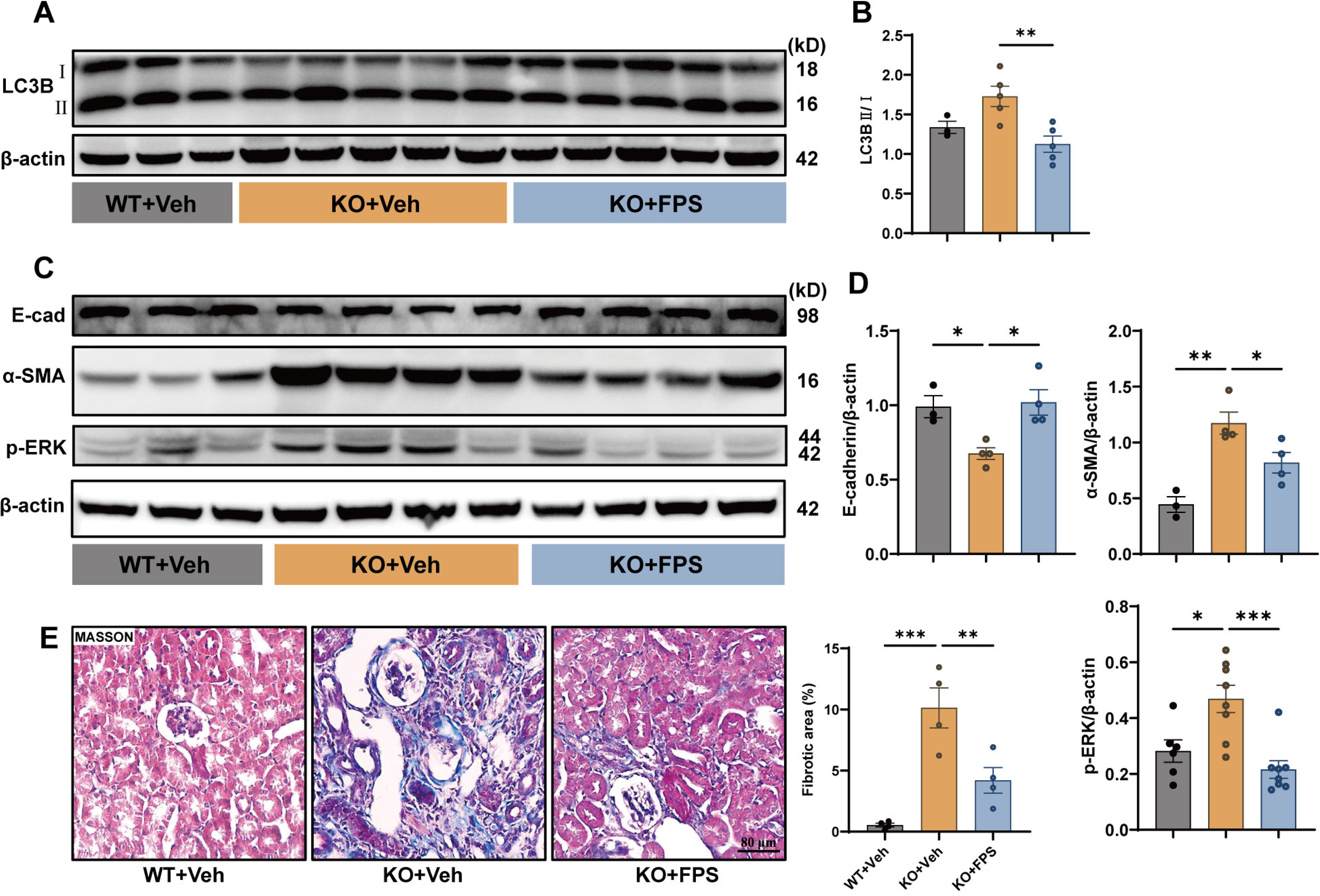
Pharmacological inhibition of RAGE mitigates renal autophagy and fibrosis in UOX^-/-^ mice. (A) Protein levels of LC3B in mouse kidney detected by WB analysis and its semi-quantification (B). (C) Protein levels of E-cadherin, α-SMA and p-ERK in mouse kidney detected by WB analysis and its semi-quantification (D). (E) Renal fibrosis shown by Masson staining. Data are presented as mean ± S.E.M (n = 3∼5 for A-B, n = 3∼4 for C-D, n = 4 for E). Scale bar = 80 μm. \**P* < 0.05, \*\**P* < 0.01.

### 9. Ferroptosis and RAGE upregulation in renal tissues of patients with hyperuricemia-related kidney disease

A preliminary exploration of renal pathology in patients with hyperuricemia-related kidney disease was conducted using para-carcinoma tissues from nephrectomy as controls. Their serological profiles are listed in Table S2. We observed iron deposition and overload in the renal tissues of patients afflicted with hyperuricemia-related kidney disease compared to the control group (Fig. 9A). IHC revealed ubiquitous high expression of GPX4 in the renal tubular epithelial cells of control subjects. However, GPX4 expression was significantly reduced in the corresponding renal regions of patients with hyperuricemia-related kidney disease, suggesting the occurrence of ferroptosis (Fig. 9B). Additionally, an upregulation of RAGE was also noted (Fig. 9C). Overall, these findings are consistent with those observed in UOX-/- mice.

**Figure 9.**
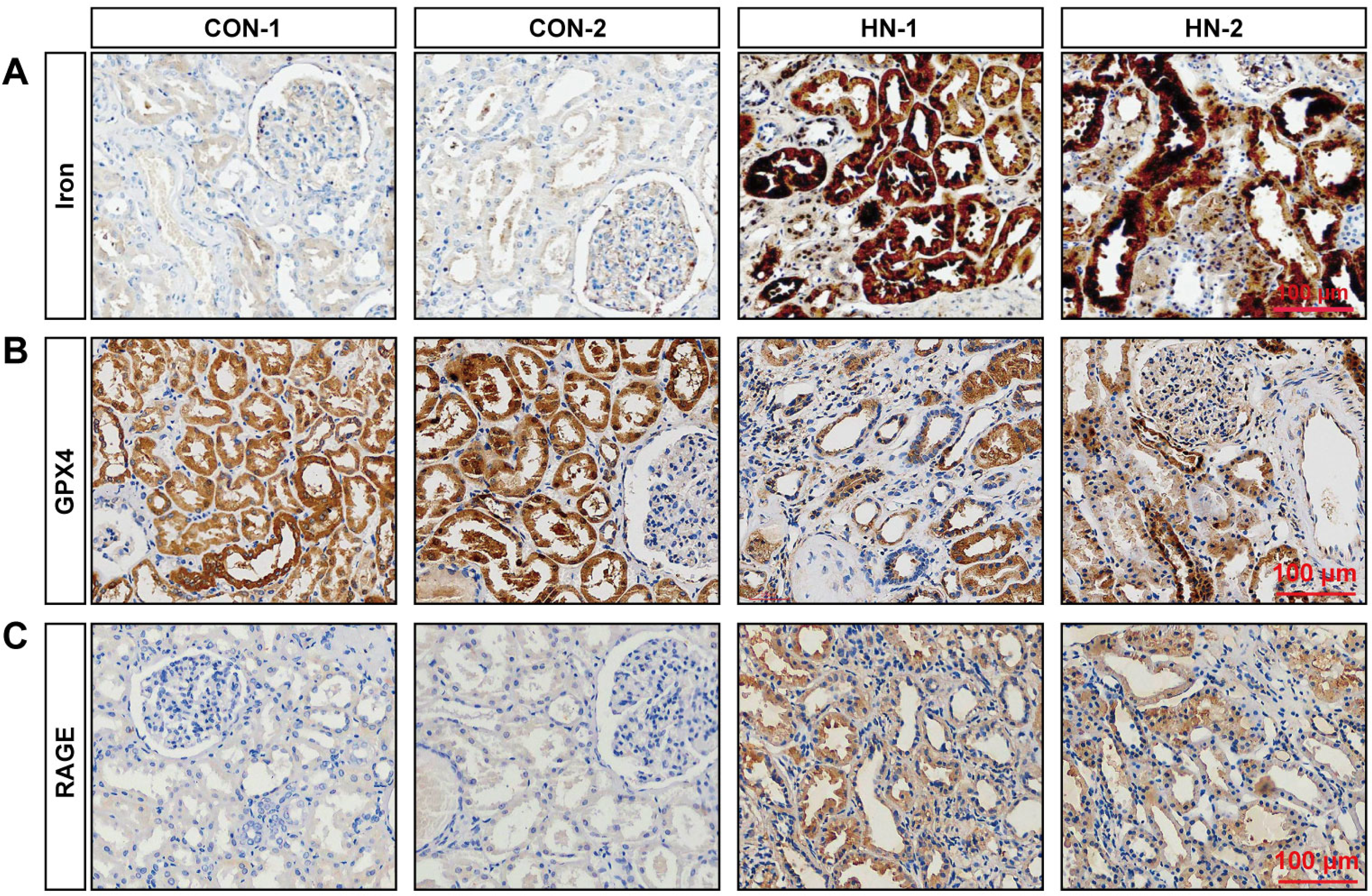
Ferroptosis and RAGE upregulation in renal tissues of patients with hyperuricemia-related kidney disease. (A) Human renal iron deposition indicated by DAB-enhanced Prussian blue staining. (B, C) Human renal GPX4 and RAGE expression indicated by IHC. CON-1 and CON-2 represent renal biopsy specimens of healthy samples from volunteer 1 and 2, respectively. HN-1 and HN-2 represent renal biopsy specimens of hyperuricemia-associated kidney diseases from patient 1 and 2, respectively. Scale bar = 100 μm.

## Discussion

Hyperuricemia is recognized as an independent risk factor for the development of renal failure ^32^, yet the unequivocal molecular underpinnings of this association remain to be fully determined. Our study contributes to the understanding of these mechanisms, presenting evidence that ferroptosis plays a critical role in mediating renal damage in HN (Fig. 10). Specifically, our data indicate that inhibiting ferroptosis markedly ameliorates kidney injury and inflammation in urate oxidase-deficient (UOX^-/-^) mice, alongside reducing lipid peroxidation, iron accumulation, and mitochondrial impairment in renal tubular epithelial cells. Furthermore, our findings highlight the role of RAGE in mediating ferroptosis-induced inflammatory injury within the kidneys of UOX^-/-^ mice. Preliminary evidence of ferroptosis and RAGE upregulation in clinical samples of hyperuricemia-associated nephropathy also supports the potential of ferroptosis and its downstream RAGE as novel therapeutic targets for HN.

**Figure 10.**
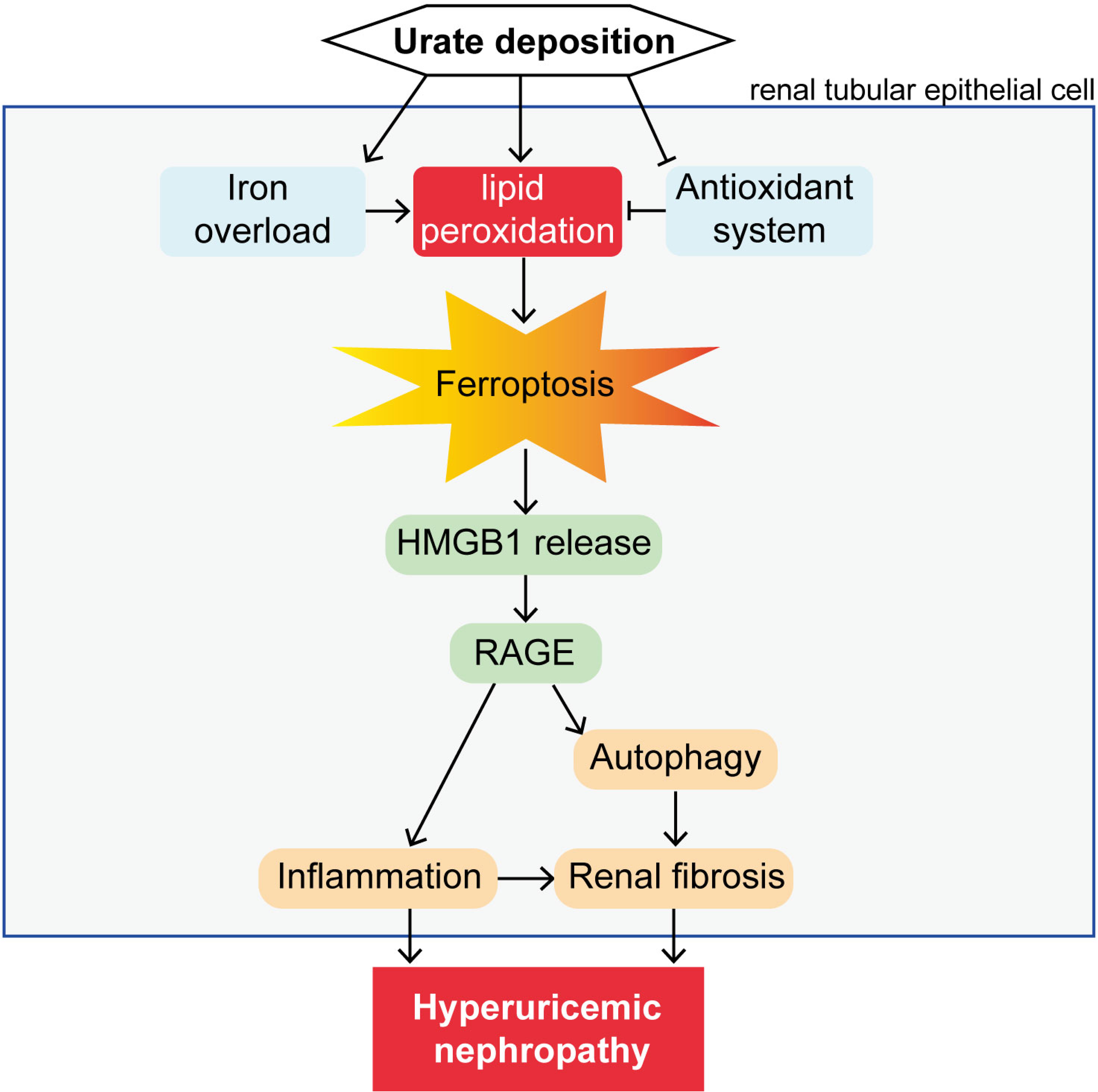
The schematic shows that ferroptosis mediates the progression of hyperuricemic nephropathy by activating RAGE signaling.

This investigation aligns with recent literature that ferroptosis in renal tubular epithelial cells promotes renal fibrosis in certain kidney injury models ^33,34^. Indeed, ferroptotic cells undergoing persistent lipid peroxidation release DAMPs, thereby continuously activating inflammatory responses, ultimately contributing to organ fibrosis. Autophagy, a cell survival mechanism under stress conditions, presents as a double-edged sword in renal diseases. While it confers a protective effect against acute renal injuries, its sustained activation in chronic renal damage encourages inflammation and fibrosis, thus impeding the recovery of renal function ^35^. Especially in the context of HN, endured autophagy activation is the key factor in inducing renal fibrosis ^36,37^. We found that in UOX^-/-^ mice, inhibition of ferroptosis attenuates renal autophagy activation, suggesting an interplay between ferroptosis and autophagy, where ferroptosis inhibition could partially improve renal inflammation and fibrosis by reducing autophagy.

In addition to ferroptosis, other forms of cell death have also been observed to play a role in HN, including apoptosis^38–40^, necroptosis^41^, and pyroptosis^42–44^, indicating a complex interplay of cell death pathways in the disease. In a mouse model of renal ischemia-reperfusion injury, Zhao et al. found that ferroptosis, necroptosis, and pyroptosis collectively constitute the main cause of acute kidney injury ^45^. However, they observed that genes related to ferroptosis were mainly expressed in renal tubular epithelial cells, while those related to necroptosis and pyroptosis were mainly expressed in macrophages, suggesting that ferroptosis may be the primary mode of cell damage for renal tubular epithelial cells. In HN, the predominant cell death pathway and the interactions among different cell death pathways require further clarification.

Numerous studies have consistently indicated that RAGE activation triggers autophagy ^20,46–48^. Given the reliance of ferroptosis on autophagy, it seemingly suggests the potential of RAGE to induce ferroptosis. Indeed, RAGE inhibition ameliorates hepatic injury in alcoholic liver disease by mitigating hepatic ferroptosis^21^. However, our investigation yielded negative results in the kidney. We discovered that RAGE inhibition improved renal outcome without affecting iron deposition or ferroptosis. This might be attributed to the inherently low basal expression levels of RAGE in renal tubular epithelial cells under normal physiological conditions. It is plausible that upregulation of RAGE occurs gradually after prolonged exposure to uric acid. By that point, ferroptosis might have already been initiated through distinct signaling pathways. Certainly, this necessitates a comprehensive temporal analysis of the sequential activation of RAGE and ferroptosis. The pivotal role of RAGE/ERK signaling in facilitating the epithelial-mesenchymal transition in renal tubular epithelial cells has been reported ^31^. Our *in vitro* and *in vivo* results also indicated an explicit suppressive effect of RAGE inhibition on ERK. Wen et al. ^28^ previously established that the HMGB1/RAGE pathway mediates the inflammatory response triggered by ferroptosis. We observed a significant reduction in elevated HMGB1 levels in the supernatant of HK2 cells and the serum of mice when ferroptosis was inhibited, which imply that HMGB1 may mediate the activation of RAGE signaling in ferroptosis in the context of HN. In summary, RAGE mediates the renal inflammatory damage caused by ferroptosis in HN.

## Conclusions

In conclusion, this study underscores the significant role of ferroptosis in hyperuricemic nephropathy. Ferroptosis mediates renal injury, inflammation and fibrosis by activating RAGE signaling in hyperuricemic nephropathy. Targeting ferroptosis and related RAGE signaling may provide novel therapeutic strategies against hyperuricemic nephropathy.

## Funding/Acknowledgement

This work was supported by grants from the National Natural Science Foundation of China (82370895, 82260163), the Natural Science Foundation of Fujian Province (2020J01018), the Joint Innovation Project of Industry-University-Research of Fujian Province (2022Y4007), and the Gout Research Foundation of Japan (Japan, 2022).

## Data Availability statement

The data used to support the findings of this study are available from the corresponding author upon request.

## Conflict of Interest

The authors declare that there are no conflicts of interest regarding this research.

## Figure legend

**Figure S1.**
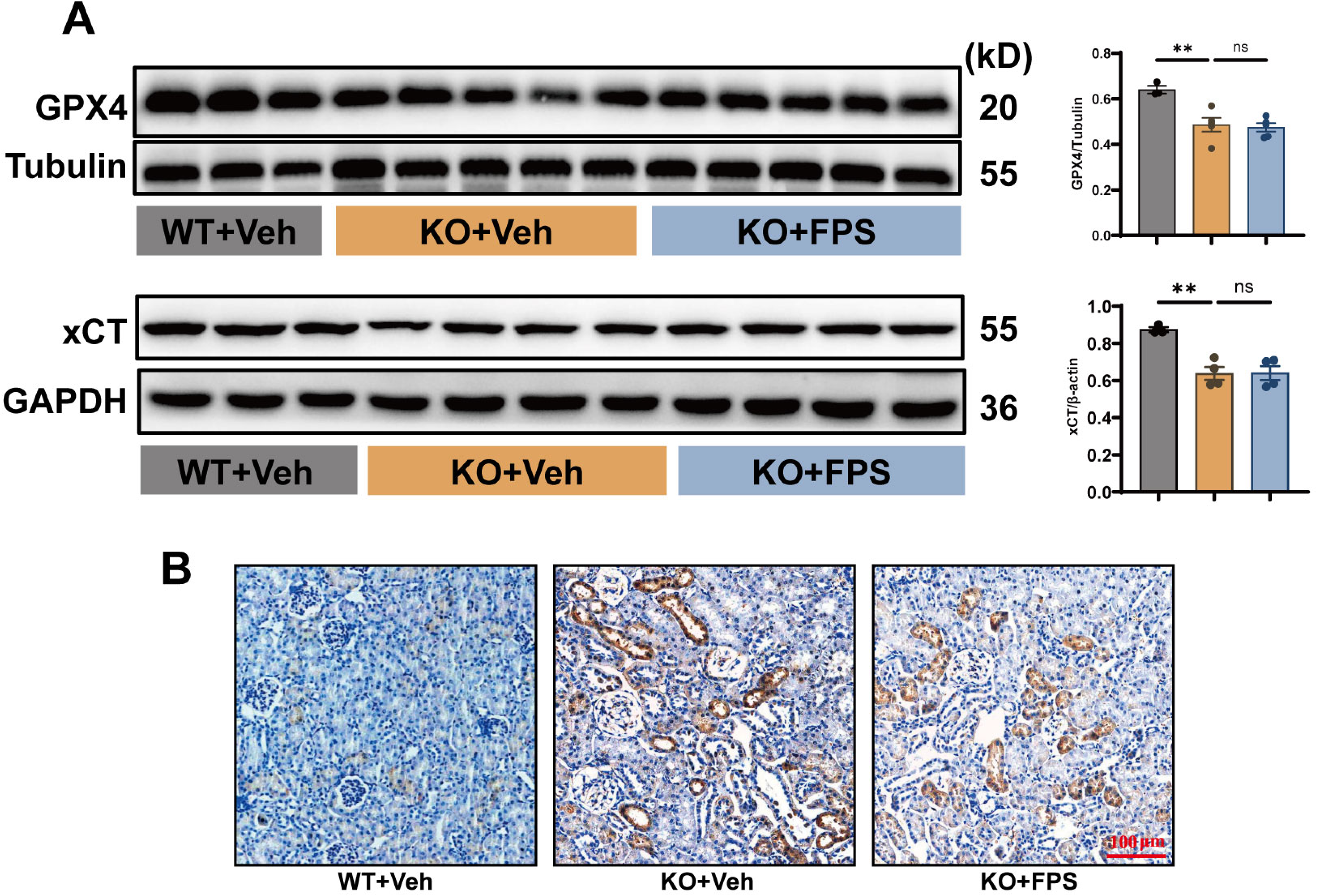
Pharmacological inhibition of RAGE alleviates renal iron deposition. (A) Protein levels of GPX4 and xCT in mouse kidney detected by WB analysis and their semi-quantification. Tubulin and GAPDH were served as internal control, respectively. (B) Iron deposition shown by DAB-enhanced Prussian blue staining. Data are presented as mean ± S.E.M (n = 3∼5). Scale bar = 100 μm.

**Figure S2.**
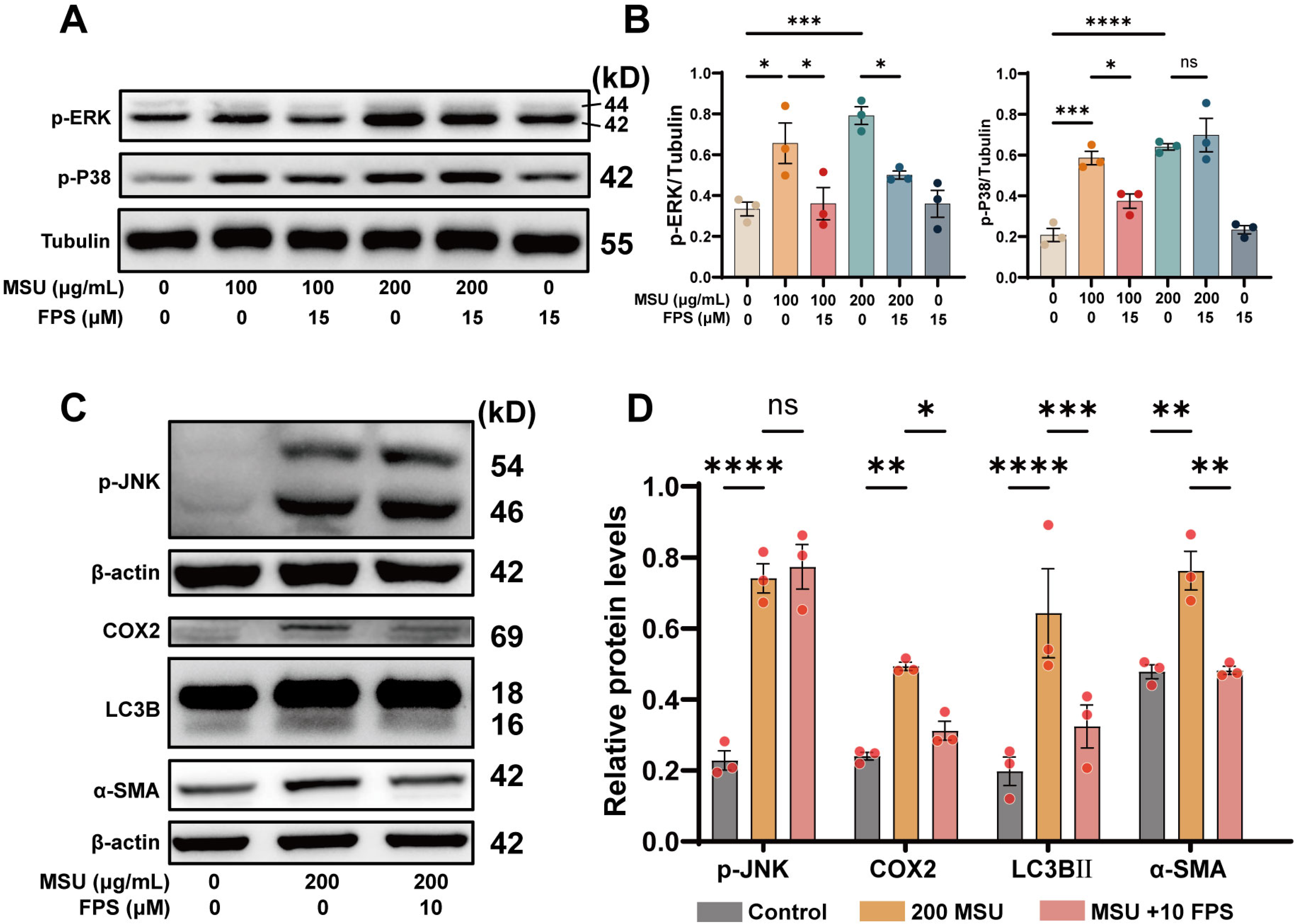
HK2 cells treated with FPS-ZM1. (A) The protein levels of p-ERK and p-P38 MAPK assessed by WB analysis and their semi-quantification (B). (C) The protein levels of p-JNK, COX2, LC3B, α-SMA assessed by WB analysis and their semi-quantification (D). Tubulin or β-actin served as internal reference proteins. Data are presented as mean ± S.E.M (n = 3). ns, not significant, \**P* < 0.05, \*\**P* < 0.01, \*\*\**P* < 0.001, \*\*\*\**P* < 0.0001.

---

**Table S1.**
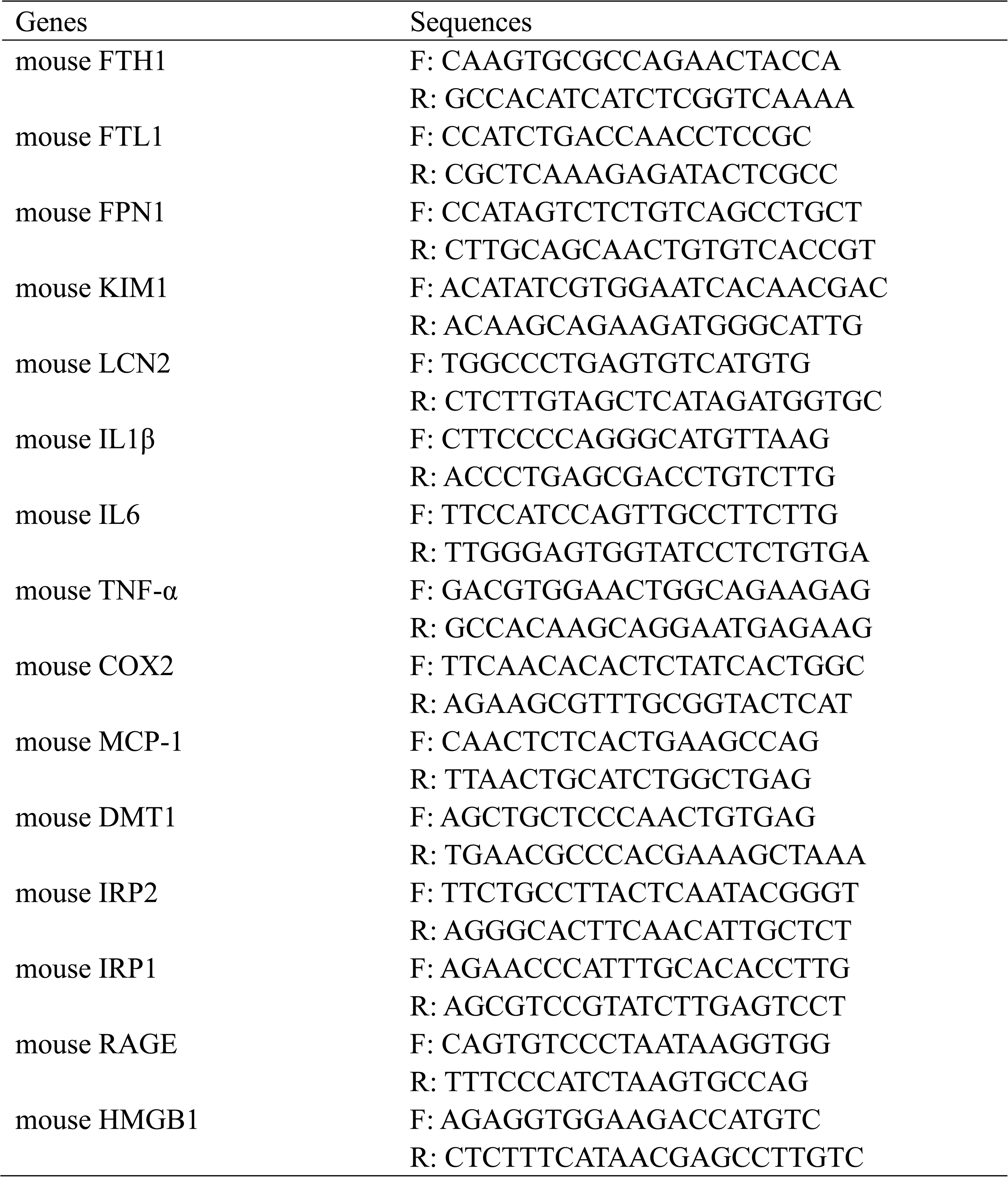
primer sequences.

**Table S2.**
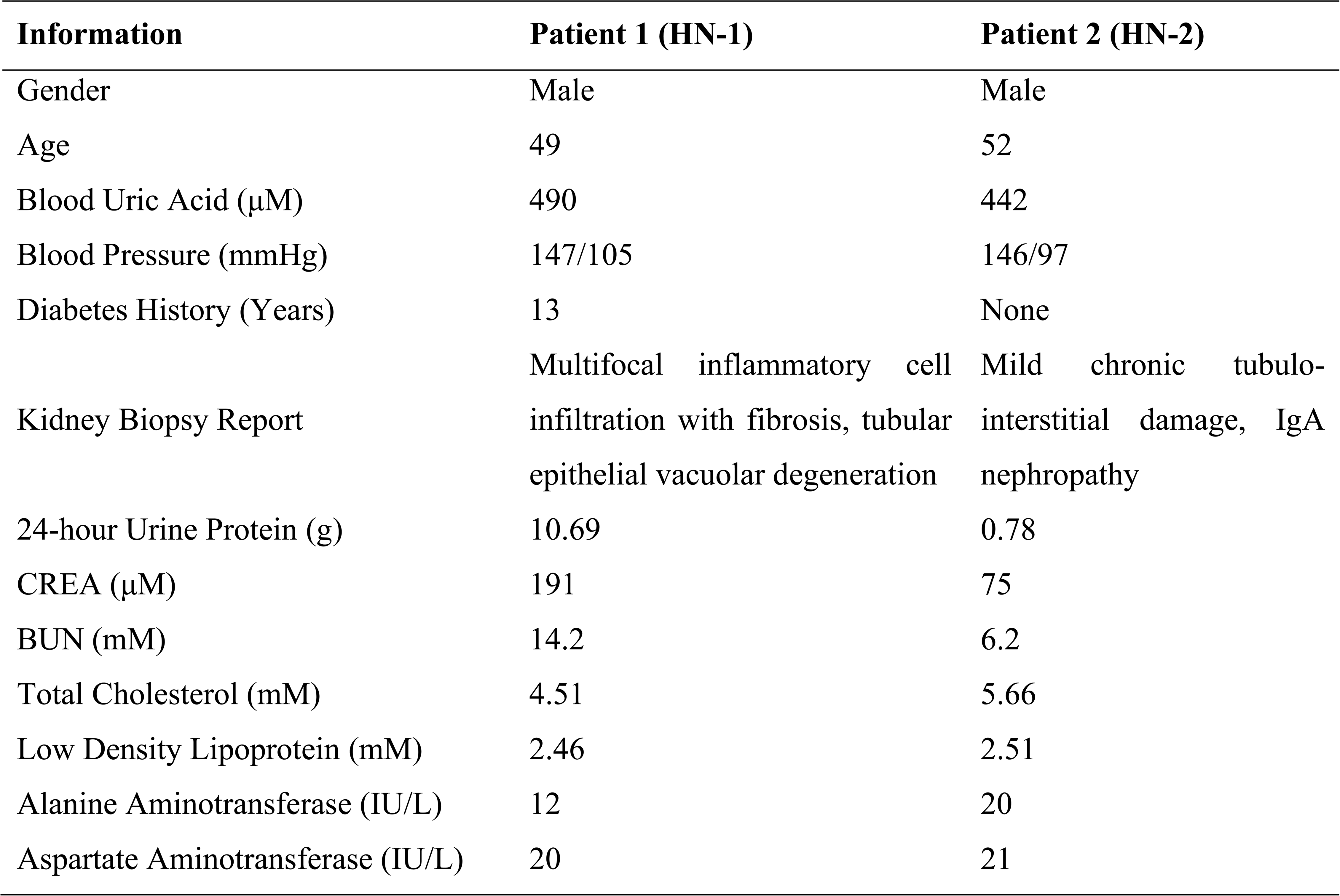
Basic Clinical Information of Patients with Hyperuricemia-Related Kidney Disease.

| Information | Patient 1 (HN-1) | Patient 2 (HN-2) |
| --- | --- | --- |
| Gender | Male | Male |
| Age | 49 | 52 |
| Blood Uric Acid (μM) | 490 | 442 |
| Blood Pressure (mmHg) | 147/105 | 146/97 |
| Diabetes History (Years) | 13 | None |
| Kidney Biopsy Report | Multifocal inflammatory cell infiltration with fibrosis, tubular epithelial vacuolar degeneration | Mild chronic tubulointerstitial damage, IgA nephropathy |
| 24-hour Urine Protein (g) | 10.69 | 0.78 |
| CREA (μM) | 191 | 75 |
| BUN (mM) | 14.2 | 6.2 |
| Total Cholesterol (mM) | 4.51 | 5.66 |
| Low Density Lipoprotein (mM) | 2.46 | 2.51 |
| Alanine Aminotransferase (IU/L) | 12 | 20 |
| Aspartate Aminotransferase (IU/L) | 20 | 21 |

## Notes

### Competing Interest Statement

The authors have declared no competing interest.

